# Chlorpyrifos degradation by combined solar photo-Fenton and bacterial metabolism and simultaneous toxicity analysis in zebrafish (*Danio rerio*)

**DOI:** 10.1101/2022.01.13.476273

**Authors:** Tanmaya Nayak, Arpan Ghosh, Sourav Das, Tapan Kumar Adhya, Paritosh Patel, Swabhiman Mohanty, Mahwish Umar, Biswadeep Das, Suraj K. Tripathy, Vishakha Raina

## Abstract

Chlorpyrifos (CP) is a widely used insecticide that has been used extensively, contributing towards a negative impact on public health concerns and associated ecosystems. Bioremediation is one of the key biological methods used for reducing these environmental toxicants. The present study examined the effectiveness of a combined process including solar photo-Fenton process followed by bacterial degradation using *Ochrobactrum* sp. CPD-03 for effective CP degradation in wastewater. Results showed that solar photo-Fenton treatment had CP degradation efficiency of ~42% in 4 h with a final degradation efficiency of ~92% in 96 h upon combined bacterial degradation. Simultaneous survivability of zebrafish (*Danio rerio*) was also studied during CP degradation. Compared to control, adult zebrafishes showed increased survivability following the addition of CPD-03 in water resulting a reduced CP concentration. CP toxicity in wastewater had caused acetylcholinesterase inhibition in zebrafish; however, this inhibition is due to absence of CP degrading bacteria. Therefore, a combined approach would influence for regulating CP degradation in wastewater along with simultaneous survival of *Danio rerio*.

## 1. Introduction

Organophosphate (OP) pesticides are a category of xenobiotic chemicals used against vectors of disease, and eye pressure relief medicines (Eaton et al., 2008). In recent decades, the use of OPs has been increased, especially in agricultural sectors, while the effects of these chemical compounds have had an adverse effect on public health (Sogorb et al., 2004). Earlier research have shown that OPs inhibit acetylcholinesterase (AChE), which inhibits the mechanism of a major neurotransmitter acetylcholine (ACh) in the central and peripheral nervous systems followed by subsequent accumulation of ACh binding to target receptors, leading to prolonged muscle contraction and eventually death through stimulation of target cells (Colovic et al., 2013; Mandour, 2013).

Chlorpyrifos (CP; *O,O*-diethyl *O*-3,5,6-trichloropyridin-2-yl phosphorothioate) belongs to OP family, act as a major pesticide used in plant pest control, domestic pests and turf pests. Toxic effects of CP has been recorded for multiple adverse health effects such as dizziness, confusion, heart rate, respiratory failure (Kavitha and Rao, 2008). In recent years, many scientists have tried to bridge the gap between CP exposure and its association towards neurodevelopmental disorders in humans and animals. CP undergoes biotransformation in which the P=S sulfur is subbed by oxygen to make chlorpyrifos-oxone (CPF-oxon) liable for AChE hindrance (Kharabsheh et al., 2017).

Water pollution has become a global problem over the last few decades. Chemical discharges in marine habitats are extremely diverse and complex, putting this ecosystem at the forefront of ecotoxicological concern as to avoid the detrimental effects of toxic molecules, such as insecticides, from natural habitat contamination (Albuquerque et al., 2016; Vieira et al., 2016). It pollutes the oceanic bodies, from point sources beginning from sewage or modern effluents and from diffuse sources, for example, rural and homegrown exercises, along these lines risking sea-going life (Akcha et al., 2012). Various studies have also reported that CP exposures have caused harmful effects on aquatic bodies and associated aquatic organisms and these include nephrotoxicity, oxidative stress, genotoxic and mutagenic effects, changes in swimming efficiency, and cell growth effects (Ali et al., 2009).

Zebrafish (*Danio rerio*), a tropical freshwater cyprinide, has as of late become an outstanding model for research in toxicology, molecular biology and vertebral evolution. It is now commonly used in numerous research areas as it holds unmatched advantages such as easy access, low maintenance costs and laboratory breeding (Lawrence, 2007). Investigation of its whole genome and the examination of quality articulation designs in an assortment of conditions uncovered its serious extent of human comparability in essential hereditary, formative and physiological cycles as 70% of protein-coding human genes are associated with zebrafish genes and 84% of human-associated genes are derived with zebrafish (Howe et al., 2013).

The fundamental issue is that pesticide-sullied water isn’t viable with conventional organic treatment because of its harmfulness to the microorganisms engaged with these cycles (van der Werf, 1996). Several microbes were isolated from OP-contaminated soil and water samples that successfully degraded CP over a period of ~30 days at different concentrations (Das and Adhya, 2015; Lakshmi et al., 2008; Lakshmi et al., 2009; Li et al., 2008). In addition, advanced oxidation processes (AOPs) are among the most frequently used industrial effluent treatment methods for water contaminated by organic compounds with high chemical stability and low biodegradability (Budarz et al., 2017). AOPs have been recognized as an substitute to waste management (Klamerth et al., 2010) and industrial wastewater pretreatment (Sirtori et al., 2009). AOPs are understood to endorse the degradation of organic compounds through the mechanism of hydroxyl radicals (·OH) formation in the primary stages. The photo-Fenton cycle has gained attention among the AOPs due to the possibility of using sunlight as a source solar energy, significantly reducing energy demand (Pintor et al., 2011; Zapata et al., 2010). These advantages include the use of low to moderate levels of chemicals, which in favorable circumstances will completely degrade the organic components of aqueous effluents; the use of simple reactors and the prospect of reuse of iron (Trovó et al., 2013).

This study aims to show the efficacy of a combined two-stage cycle (photo-Fenton followed by bacterial degradation) to treat CP. This also validates the extent that bacterial culture *Ochrobactrum* sp. CPD-03 serves as an effective bioremediator to reduce the CP toxicity for zebrafish in the same ecosystem.

## 2. Materials & methods

### 2.1. Test chemical and zebrafish maintenance (egg production, embryo exposure, morphological observation)

Pure standards of CP (analytical grade) used in this study was purchased from Sigma-Aldrich (St.Louis, USA). Stock solution of CP (1000 μg mL^−1^) was prepared upon dissolving accurate amounts of pure standards in dimethilsulfoxide (DMSO) (molecular grade). Acetonitrile, DMSO and HPLC grade water were purchased from Merck (Darmstadt, Germany). Zebrafish (wild-type) were obtained from the in-house Zebrafish laboratory facility (School of Biotechnology, KIIT University). Zebrafish embryos were maintained in the embryonic media containing NaCl, 0.29 g mL^−1^; KCl, 0.012 g mL^−1^; CaCl_2_•2H_2_O, 0.048 g mL^−1^ and MgSO_4_•7H_2_O, 0.081 g mL^−1^. This embryonic media was prepared in 1000x stock concentration followed by autoclaved and stored in room temperature. Zebrafish maintenance was carried out in a dish maintaining system supplied by Aquaneering (San Diego, USA). The water parameters were as follows: temperature: 27±0.5 °C, Conductivity: 744 μs, hardness: 379 mgL-1 CaCO_3_ (21.30 dH), pH: 7.5 ± 0.25, dissolved oxygen: 10.5±0.5 mg L^−1^ O_2_ (95% saturation) and 12:12 dark: light photoperiod. Fishes were fed with dried bloodworm available in the market. The breeding setup was in a breeding tank, supplied by Aquaneering, in fish water with 1:2 male: female ratio. All procedures were approved by the Research and Development Committee of the Institutional Animal Care and Use Committee at KIIT University and carried out under a license from the local government in compliance with the institutional guidelines.

### 2.2. Experimental setup

#### photo-Fenton degradation

Photo-Fenton experiments were carried out in 500 mL glass beaker containing minimal salts media (MSM) (Nayak et al., 2020) supplemented with CP (100 mg L^−1^) under continuous and controlled agitation (200 rpm). The beaker was kept in solar light (with an illuminance ≈100,000 ± 20000 lx). During the experiment, the device temperature was controlled by a digital thermometer and held at 35 ± 3 °C. Furthermore, 100 mg L^−1^ hydrogen peroxide (reagent grade 35% w/v, Merck) was added. Samples were collected at every 1 hour intervals to monitor the degradation process.

#### Bacterial degradation

The MSM media containing residual CP after the photo-Fenton process was filter sterilized using a 0.22 μm membrane filter (Merck, Germany). To this media, *Ochrobactrum* sp. CPD-03 (1.6×10^5^ cfu ml^−1^) was added. Another set up was operated without the bacteria. In addition to this, a model organism zebrafish (*Danio rerio*) was also added to the setup to monitor the effect of pesticides with the morphological and behavioral studies. To test the bacterial count, 100 μL of the samples collected were diluted in regular sterile saline (0.9% NaCl) and 100 μL of the correct dilution was placed on LB agar plates. The plates were incubated overnight at 37 °C.

### 2.3. Chlorpyrifos exposure

Analytical grade chlorpyrifos (CP, 99.9% purity, Sigma-Aldrich, USA) was used in this study. CP stock solutions were prepared in DMSO (0.9%) and this mixture was added to MSM, ensuring solubility. To the sterile MSM (sterilized with 0.22 μm membrane filter as mentioned above) obtained after photo-fenton process, acclimated fishes (n=9) were placed into a glass beaker containing 50 ml of treatment, these treatments were performed in periods of 96 h over an interval of 24 h. After the procedure, each fish was removed from the system, sacrificed and deposited in a freezer of −20^0^C before acetylcholinesterase was analyzed. The MSM content of the beaker was subjected for HPLC analysis.

### 2.4. Bacterial culture preparation

*Ochrobactrum* sp. CPD-03 was previously isolated in the lab and have been studied for CP degradation in aqueous media. Pure cultures of bacterial strain CPD-03 were cryopreserved in 25% glycerol at −80^0^C. The bacterial strain was revived and inoculated in a 100 ml Erlenmeyer flask with 20 ml sterile MSM supplemented with CP (100 mg L^−1^) before each set of experiments. Flask was incubated at 30 ± 2 °C with 120 rpm in a shaker incubator. The bacterial cells were harvested after 44 - 48h at 4500 rpm for 5 min followed by washing with sterile saline (0.9% NaCl). The washed bacterial cells were resuspended in sterile MSM to obtain a cell density of 1-1.5×10^8^ cells ml^−1^.

### 2.5. Biochemical assays: apoptosis, acetylcholinesterase activity, glutathione S-Transferase activity

A preliminary analysis was performed for the morphological changes in zebrafish embryo upon CP (100 mg L^−1^) exposure along with CPD-03 (1.6×10^5^ cfu ml^−1^) for 96 h. Moreover, apoptosis due to CP exposure at various concentration was analysed by staining the zebrafish treated embryos/larvae (7dpf) with acridine orange stain. The stock stain solution of 3mg/ml was prepared in 10X PBS. The working stain solution (7μg/ml) was prepared by diluting the stock with distilled water. Embryo/larvae (n=9) from all the concentrations was removed and rinsed with embryonic medium. The larvae were then transferred to 24 well-plates containing 1ml of embryonic medium with 20μl of acridine orange stain and incubated for 20 minutes in dark. The apoptotic cells were presented and photographs taken in the green channel of the inverted fluorescent microscope.

Previous report have shown that CP is toxic to the fish by inhibiting brain acetylcholinesterase (AChE) activity (Tiwari et al., 2019). In this study, frozen heads of zebrafish (control and treatment exposure groups) were thawed and homogenized in 0.1M sodium phosphate (NaPO_4_; pH 7). Tissues were homogenized for approximately 45 s with a polytron homogenizer (Powergen, Fisher Scientific, and Hanover Park, IL). Throughout homogenization, tissues stayed on ice. The homogeneous material was then centrifuged at 12,000 g for 5 min and the supernatant was transferred for biochemical processing, specifically for total protein and AChE activity.

Total protein was determined by Coomassie blue method using Bradford dye (Sigma-Aldrich, St. Louis, MO, USA) and absorbance was measured at 595 nm. Bovine Serum Albumin (BSA) (Sigma-Aldrich, St. Louis, MO, USA) was used as the standard. Around 20 ml of sample supernatant were used from each sample to measure protein content using the Bradford method (Bradford, 1976). The quantity of absorption was proportional to the protein present.

Homogenized sample supernatants were then tested for the activity of acetylcholinesterase (Ellman et al., 1961) using a slight modification. In short, the activity of acetylcholinesterase from the supernatant was calculated using 99% of acetylthiocholine iodide (Sigma-Aldrich, St. Louis, MO, USA) as the substrate in a 2 ml reaction medium consisting of 0.25mM 5, 5-dithio-bis (2-nitrobenzoic acid) or DTNB (Sigma-Aldrich, St Louis, MO, USA), 0.1 M sodium phosphate buffer (Sigma-Aldrich, St. Louis, MO, USA) at pH 7.5, and 0.001 M acetylthiocholine iodide. Absorbance was read at 412 nm. The activity of enzymes in the supernatant was standardized by total protein as determined by the Bradford method described above. The AChE activity was replicated three times and repeated at least two times for each treatment.

Glutathione-S-transferase activity was determined by using EZAssayTM GST Activity Estimation kit (HiMedia, India). The enzyme assessment was conducted based on protocol provided along with the kit. The reaction mixture was prepared in 1ml quartz cuvette and assay buffer (930μl) was used to set the spectrometer to zero. Tissue homogenate was prepared by homogenizing embryos/larvae (n=9) in cold diluted sample buffer using motar pestle. The reaction mixture consisted of 6-10 μl 1-chloro-2,4-dinitrobenzene (CDNB) as (substrate), 8-10μl glutathione (GSH), 3-6μl Glutathione S-transferases (GST) standard and 50μl of sample. The increase in absorbance was measured in duplicates at 340 nm at every 1 min time interval using spectrometer.

### 2.6. Sample extraction & HPLC analysis

The MSM media after photo-Fenton treatment and bacterial degradation process, was harvested at 7500 rpm for 10 min for obtaining a cell-free media and extracted with organic solvents. The supernatant was pulled out for extraction with an equal volume of ethyl acetate and this step was repeated twice. The organic layer thus obtained was aspirated, pooled and dehydrated over a column of anhydrous sodium sulfate, evaporated at 45±2°C using rotavapor (Eyela, Japan), reconstituted in methanol (HPLC Grade) and filtered through 0.22 μm membrane filter. Residual CP and its metabolite 3,5,6-trichloro-2-pyridino (TCP) was quantified using a reverse-phase C18 column (ZORBAX Eclipse Plus, Agilent Technologies, USA) fitted with Agilent Technologies 1260 Infinity HPLC system equipped with a binary pump, UV/VIS detector, thermostat column compartment (TCC) column oven and auto sampler, with array detection based on peak area and retention time of the pure standard. The mobile phase used for the analysis contained a mixture of methanol and water in a ratio of 85:15 at a flow rate of 1.2 mL min^−1^. Water (HPLC grade) was acidified to pH 3.2 using orthophosphoric acid. Injection volume was 20 μL. The recoveries of OPs added at concentrations of 1, 2, 5, 10, 50 mg/L, ranged from 97.0% to 100.0%. The linear range of the calibration curves was obtained from 1 to 50 mg/L.

### 2.7. Data analysis

All statistical analyses were performed by GraphPad Prism and assessed by ANOVA. The data were expressed as mean ± SEM, and p<0.05 or below was considered for significant difference.

## 3. Results & discussion

### 3.1 Photo-Fenton treatment of the chlorpyrifos

Several photo-assisted blank experiments are conducted under conditions such as photocatalytic assays to ensure consistent results and no other effects of H_2_O_2_. Earlier studies have illustrated the oxidation process utilizing hydrogen peroxide (H_2_O_2_) as oxidant and iron (Fe) as a catalyst in the presence of acidic (H^+^) medium (Fenton, 1894). Briefly the fenton oxidation process starts with generation of a hydroxyl free radical (·OH) (Dewil et al., 2017) and these are one of the most active oxidants which could react 106-1012 times faster than ozone depending on the substrate needs to be degraded (Hoigné, 1997; Lloyd et al., 1997). At early stages of the photo-Fenton reaction, molecules of pesticide are degraded by hydroxyl radicals, leading to organic intermediates formation. The steps involved in the Fenton reaction are described as below Equations (1)–(7) (Javaid and Qazi, 2019).

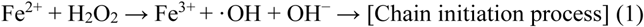

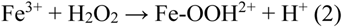

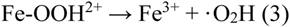

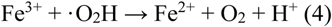

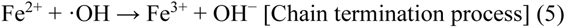

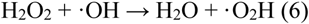

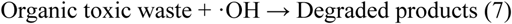

In this study, a preliminary assessment was conducted to ensure the sun-light assisted photo-Fenton approach (with ~1,20,000 lx) with and w/o H_2_O_2_ (25 mg L^−1^) to ensure CP degradation. H_2_O_2_ concentration was kept as per an earlier report (Villegas-Guzman et al., 2017). FeSO_4_ present in MSM media worked as a fenton reagent. Up to ~42% of CP degradation in the sun-light assisted photo-fenton process was achieved (Fig.1) in initial 4 h of the reaction with a degradation efficiency of 42.61%. No further CP degradation was observed afterwards, hence the reaction was stopped. As per an earlier report, this above reaction gradually reduced the radical production followed by depletion in iron and H_2_O_2_ concentration which will not allow the reaction (Malato et al., 2009). Possibly this could be the reason behind CP degradation hinderance beyond 4 h. TCP detection upon CP degradation could not be captured. This might be due to further

**Fig. 1:**
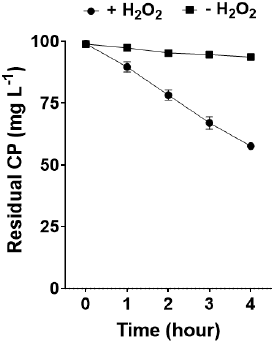
Time course CP degradation is driven by presence of H_2_O_2_. Data were represented in the mean of n=5 and analysis were performed using ANOVA with the Prism8. All were tested at the *p*<0.001*** significance level

### 3.2 Biological degradation of Chlorpyrifos and Zebrafish survivability

The survivability of zebrafish larvae upon exposure with various CP concentration (100 – 500 μg ml^−1^) and was found that higher concentration led to cell damage (Fig. 2). A control set was kept in DMSO (0.9%) as beyond this concentration would have toxic to the zebrafish (Maes et al., 2012). Higher CP concentration (<200 μg ml^−1^) has shown to negatively affect the cell morphology thereby increasing the fluorescence. Earlier research has been conducted to determine the exposure of CP on animals, specifically rats (Terry Jr et al., 2012), and zebrafish (Eddins et al., 2010), and investigated their relevance to humans. Based on these, zebrafish survivability was observed for 96 h during bacterial degradation of the residual CP from photo-Fenton process (Fig.3a). Significant differences were observed between the presence and absence of CPD-03 during residual CP exposure. As depicted in the fig. 3a, zebrafish population was reduced to ~50% in 12 h and ~100% in 24 h in the absence of CPD-03 during residual CP exposure, whereas presence of CPD-03 also has shown reduced zebrafish survivability but in a later stage. It is possibly due to the presence of CPD-03 has reduced the CP concentration which might have supported the zebrafish survival.

**Fig. 2:**
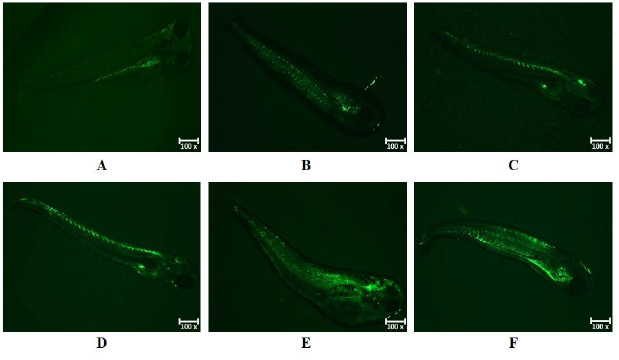
Zebrafish survivability analysis by apoptosis in presence of various concentration of CP. A-Solvent control (0.9% DMSO), B-100 μg mL^−1^, C-200 μg mL^−1^, D-300 μg mL^−1^, E-400 μg mL^−1^, F - 500 μg mL^−1^

**Fig. 3:**
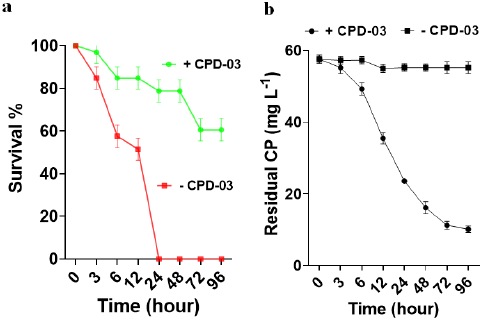
a. Zebrafish survivability in presence of *Ochrobactrum* sp. CPD-03 (1.6×10^5^ cfu ml^−1^); b: Time course zebrafish survivability in presence *Ochrobactrum* sp. CPD-03. Data were represented in the mean of n=9 and analysis were performed using ANOVA with the Prism8. All were tested at the *p*<0.005*** significance level; b. Time course CP degradation is driven by addition of *Ochrobactrum* sp. CPD-03 (1.6×10^5^ cfu ml^−1^). Data were represented in the mean of n=5 and analysis were performed using ANOVA with the Prism8. All were tested at the *p*<0.001*** significance level.

Previous reports have validated that the combined approach including photo-Fenton assisted followed by bacterial degradation have shown higher degradation efficiencies for pesticide degradation (Martín et al., 2009). In addition to this, one of our earlier studies have shown the efficiency of *Ochrobactrum* sp. CPD-03 towards CP (100 mg L^−1^) degradation in aqueous medium (Nayak et al., 2020). Moreover, the H_2_O_2_ concentration used in photo-Fenton was below sublethal concentration for microbes (Labas et al., 2008). The residual CP in photo-Fenton treatment was further treated with the presence of the bacterial strain, *Ochrobactrum* sp. CPD-03 to illustrate the efficiency of this combined degradation approach. In this study, CP degradation was found positively regulated with the presence of CPD-03 as depicted in the fig. 3b. This could be possible that, the resulting TCP would have been degraded by CPD-03 and hence it could not get detected in this time course. The CP degradation efficiency of 90% was found in case of CPD-03 addition in 96 h.

### 3.4 Acetylcholinesterase (AChE) activity

Acetylcholinesterase (AChE) activities varied in different time during CP exposure as AChE is the most common targets for OP-based chemical compounds. AChE is inactivated by OP chemicals and prevents ACh hydrolysis. Most of the OP studies used AChE as a biomarker for OP toxicity and exposure. The AChE test has proved to be considerably reliable method for detecting the level of CP toxicity in mammals in particular. Earlier studies were reported that the higher concentration of pesticides has decreased the AChE activity (Mishra and Devi, 2014; Xing et al., 2014). Our study has shown significant decrease in AChE with increased CP concentration (Fig.4a). In this study, AChE activity was significantly increased in the zebrafish with the residual CP exposure in the absence of CPD-03 (Fig.4b). AChE activity found considerably decreased in the presence of CPD-03 as the residual CP was degraded by the bacterial metabolism (Fig.4c). It might be due to the CP degradation occurred by CPD-03 which had depleted the concentration of CP and resulted a significant less exposure to the zebrafish (Fig.4d).

**Fig.4 (a-d):**
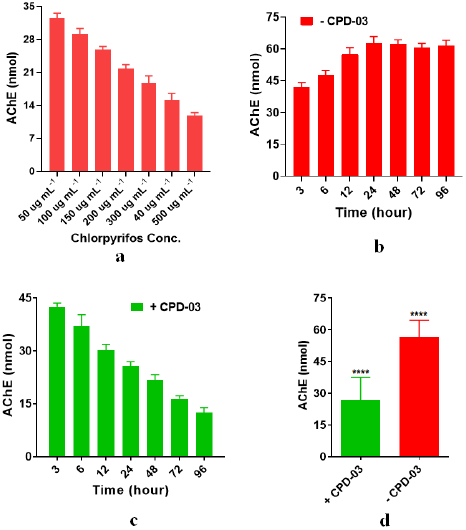
a. AChE activity upon increased CP concentration; b-d. AChE activity in presence *Ochrobactrum* sp. CPD-03 (1.6×10^5^ cfu ml^−1^); Data were represented in the mean of n=5 (system set-up) and analysis were performed using ANOVA with the Prism8. All were tested at the *p*<0.005*** significance level

### 3.5 Glutathione-S-Transferase Assay

Glutathione S-transferases (GST) are among the main antioxidant enzymes and the primary antioxidant markers (Yang and Lee, 2015). Nonetheless, GST plays a central role by detoxifying xenobiotic compounds (Higgins and Hayes, 2011) and glutathione-conjugated endogenous compounds (Domingues et al., 2010). In this study, in the absence of CPD-03, the oxidative stress enzyme GST was inhibited at later times (Fig. 5). This inhibition may be due to a GST deficiency in response to a significant increase in stress due to the presence of CP. Significant results were obtained in previous reports (Oliveira et al., 2013) and was found in embryos of zebrafish when xenobiotic compounds were exposed for 96 h without any degradation carrier (Melo et al., 2015). The presence of CPD-03 has shown lesser GST activity over a period of time as the CP degradation occurred by CPD-03.

**Fig.5:**
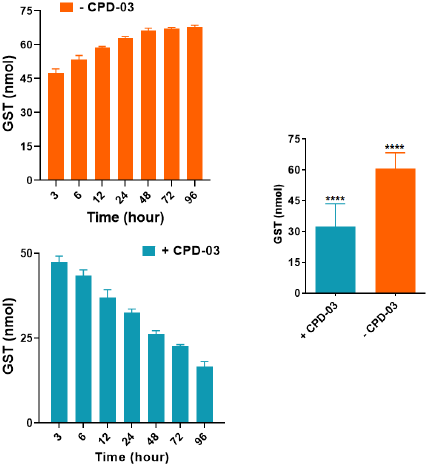
Time course GST activity in presence *Ochrobactrum* sp. CPD-03 (1.6×10^5^ cfu ml^−1^); Data were represented in the mean of n=5 (system set-up) and analysis were performed using ANOVA with the Prism8. All were tested at the *p*<0.005*** significance level

## 4. Conclusion

Sublethal exposure to CP can be concluded to lead to oxidative stress in zebrafish at 96 h. Results found using a biodegradability assessment method that is simple, quick and economical using *Ochrobactrum* sp. CPD-03 has allowed us to determine the optimal strength of photo-Fenton treatment for hazardous, non-biodegradable wastewater (CP-containing) treatment. The combination of photo-Fenton treatment accompanied by biological degradation proved to be an effective treatment for rapid degradation of pesticides (96 h) in wastewater (containing ~60-70 mg L^−1^ of CP) without causing much damage to the aquatic organisms. In zebrafish, The primary cause of pesticide poisoning may be respiratory failure, i.e. muscle inability to control the amount of water (and therefore oxygen) in the gills, resulting in reduced blood oxygenation, leading to hypoxia-induced death. This study reports the use of brain AChE assay to assess the effects of CP in the presence and absence of *Ochrobactrum* sp. CPD-03 in adult zebrafish. The results indicated significantly lesser inhibition of brain AChE activity in all CPD-03 treated groups compared to untreated water groups. Additional studies should include muscle inhibition assessments of AChE, particularly those that control the operculum. Furthermore, since CP intoxication has shown a significant reduction in the activity of glutathione s-transferase, these enzymes should be measured to better understand. As far as we know, most of the literatures have studied the photo-fenton and bacterial degradation of pesticides differently. This study would be the first report in a combined approach including photo-fenton followed by bacterial degradation for the removal of parent compounds like CP. Future studies should focus on to enhance the activity of this combined approach by the application of bioreactors for efficient removal of pesticides. Nevertheless, evaluation of this technology should take into account for the higher concentrations of pesticides and how photocatalytic and biological oxidation are influenced by this factor.

## Authors’ contributions

TN & VR designed the research and outline. TN, AG & SD performed the experiments and generated the result. SKT supported photo-fenton data analysis. PP & MU performed the zebrafish maintenance and experiments. BD reviewed the zebrafish results. TN wrote the manuscript. TKA & VR critically reviewed the manuscript with appropriate technical inputs. All authors have read and approved the final manuscript.

## Declaration of interests

The authors declare that they have no known competing financial interests or personal relationships that could have appeared to influence the work reported in this paper.

## Acknowledgement

This research work was funded by the Department of Biotechnology (DBT), Govt. of India, New Delhi. Research grant sanction no. BT/PR7580/BCE/8/1009/2013. All authors remain highly grateful to the Director of the institute for infrastructure that enabled us to perform this work.

---

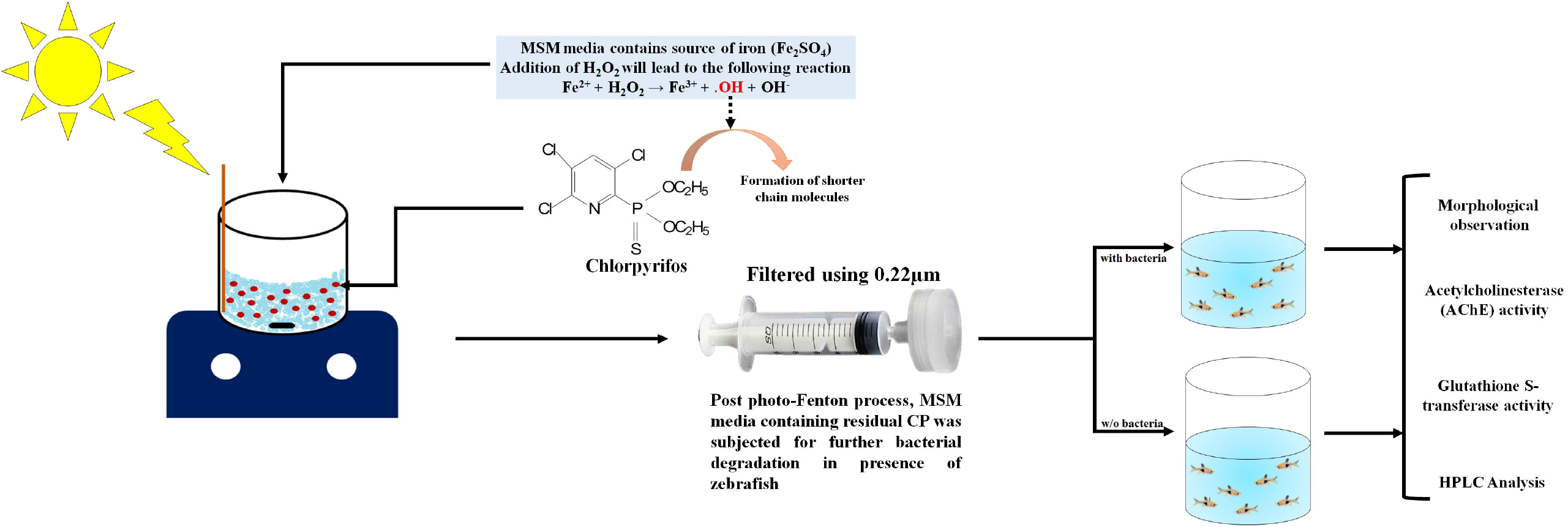

## Notes

### Competing Interest Statement

The authors have declared no competing interest.

